# Neutrophil extracellular traps induce the epithelial-mesenchymal transition: implications in post-COVID-19 fibrosis

**DOI:** 10.1101/2020.11.09.374769

**Authors:** Laura Pandolfi, Sara Bozzini, Vanessa Frangipane, Elena Percivalle, Ada De Luigi, Martina Bruna Violatto, Gianluca Lopez, Elisa Gabanti, Luca Carsana, Maura D’Amato, Monica Morosini, Mara De Amici, Manuela Nebuloni, Tommaso Fossali, Riccardo Colombo, Laura Saracino, Veronica Codullo, Massimiliano Gnecchi, Paolo Bigini, Fausto Baldanti, Daniele Lilleri, Federica Meloni

**Author notes:** Corresponding author: Laura Pandolfi, PhD, IRCCS Policlinico San Matteo Foundation, Viale Golgi 19, 27100, Pavia, Italy.

## Abstract

The release of neutrophil extracellular traps (NETs), a process termed NETosis, avoids pathogen spread but may cause tissue injury. NETs have been found in severe COVID-19 patients, but their role in disease development is still unknown. The aim of this study is to assess the capacity of NETs to drive epithelial-mesenchymal transition (EMT) of lung epithelial cells and to analyze the involvement of NETs in COVID-19.

Neutrophils activated with PMA (PMA-Neu), a stimulus known to induce NETs formation, induce both EMT and cell death in the lung epithelial cell line, A549. Notably, NETs isolated from PMA-Neu induce EMT without cell damage. Bronchoalveolar lavage fluid of severe COVID-19 patients showed high concentration of NETs. Thus, we tested in an *in vitro* alveolar model the hypothesis that virus-induced NET may drive EMT. Co-culturing A549 at air-liquid interface with alveolar macrophages, neutrophils and SARS-CoV2, we demonstrated a significant induction of the EMT in A549 together with high concentration of NETs, IL8 and IL1β, best-known inducers of NETosis. Lung tissues of COVID-19 deceased patients showed that epithelial cells are characterized by increased mesenchymal markers. These results show for the first time that NETosis plays a major role in triggering lung fibrosis in COVID-19 patients.

## Introduction

Neutrophils represent the most abundant type of white blood cells. Neutrophils protect against foreign pathogens and are considered an essential component of the innate immune system. After activation, neutrophils can react against pathogens with three major different mechanisms: i) phagocytosis; ii) release of granules (that contain proteases and ROS); iii) NETosis. This last mechanism is considered a type of programmed neutrophils cell death, which consequently releases NETs (1). NETs are webs-like structures composed by chromatin decorated with proteases, such as human neutrophils elastase (HNE) and myeloperoxidase (MPO), whose primary role is limiting the spreading of pathogens in tissues (2). Several signals can stimulate NETosis: microorganisms; pro-inflammatory cytokines, such as IL8 and IL1β; and chemicals, such as PMA (3).

In spite of their protective role in reducing pathogens diffusion in the parenchyma, several studies revealed that a persistence of NETs can amplify the primary injury therefore inducing further clinical complications (4, 5, 6). In the lung, the persistence of activated neutrophils that release NETs has been linked to several diseases, such as cystic fibrosis and acute respiratory distress syndrome (ARDS). Notably, COVID-19 patients who developed ARDS show increased NETs in the serum and, most importantly, NET release significantly correlates with the severity of the lung pathology (7). Moreover, a recent paper demonstrated that NETs are detectable in tracheal aspirate of patients with COVID-19 and that they are involved in the prothrombotic clinical manifestations of COVID-19 (8). The direct consequence of NETs persistence in the lung is the damage of epithelial and endothelial cells, driven predominantly by histones (1). Additionally, NETs have been found to drive the EMT in the context of the breast cancer thanks to their capacity to upregulate transcription factors involved in the EMT (*ZEB1* and *SNAI1*) (9).

Given these premises, the aim of this study is to assess whether NETosis may drive the EMT also in lung epithelial cells. This would represent a new crucial molecular mechanism underlying the development of inflammatory-induced lung fibrosis, thus it would also be of great importance for several pulmonary diseases, such as autoimmune microbial insults and transplant rejection. Also, given the high concentration of NETs in COVID-19 patients that develop ARDS, this study is aimed at understanding if NETs could be implicated in the induction of EMT in SARS-CoV2 pneumonia by promoting a fibrosis.

## Results

### NETosis induces the EMT in an alveolar epithelial cell line

Since NETosis is toxic in the lung (4, 6) and it induces the EMT in breast cancer cells (9), we investigated if NETs can induce the EMT in the alveolar epithelial cell line A549. We used PMA as a primary chemical effector of NETosis. Confocal microscopy confirmed that PMA leads to a very efficient release of NETs by human neutrophils (Fig. 1L-T) compared to non-activated cells (Fig. 1A-I). NETs are visible as web-like structures and, as previously reported (10, 11), they are positive for the histone H3 (Fig. 1L-N), HNE (Fig. 1O-Q) and MPO (Fig. 1R-T). Moreover, DAPI labeling (Fig. 1L, O, R) further confirms that these proteins are associated to free-DNA (Fig. 1N, Q and T).

**Figure 1.**
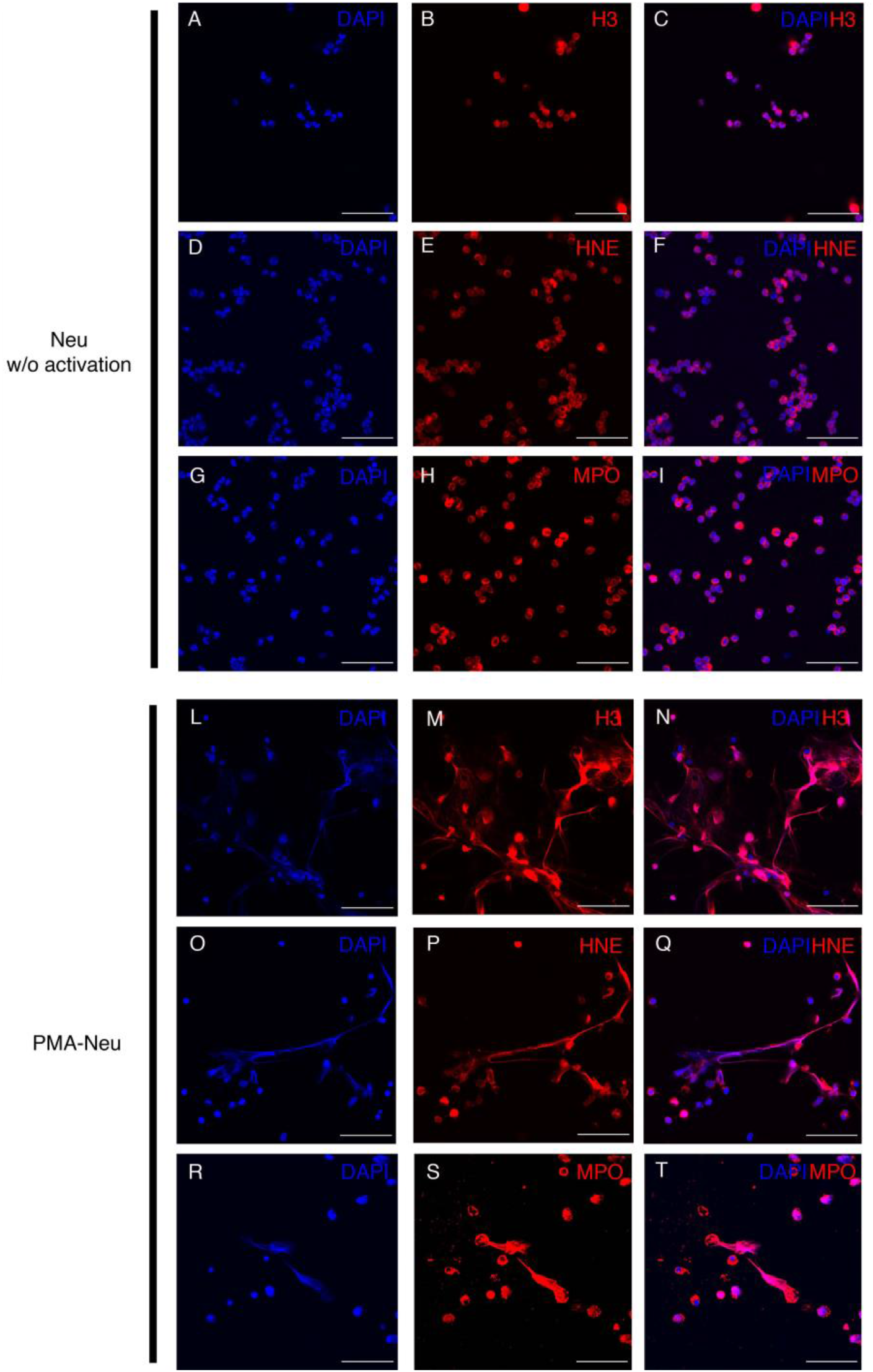
Confocal images of neutrophils activated or not towards NETosis. (A-I) Neutrophils isolated from peripheral blood of healthy donors without any treatment or (L-T) treated with 100 nM of PMA for 4 h. All samples were then labeled with (B,M) anti-H3 or (E,P) anti-HNE or (H,S) anti-MPO Ab (red signal). (A,D,G,L,O,R) DAPI (blue signal) was used to label DNA. (C,F,I,N,Q,T) Images of merged channels. Scale bar = 50 μm.

To evaluate the possibility that NETs may induce the EMT in lung epithelial cells, we cultivated A549 cells with PMA-Neu for 24 and 48 h. By microscopy we observed that A549 cells lose the epithelial morphology upon incubation with PMA-Neu (Fig. 2A) and gain a fibroblastic phenotype (Fig. 2C). These morphological changes are visible also after TGF-β treatment, the best-known effector of the EMT (Fig. 2B). To confirm that these morphological changes are related to EMT induction, we evaluated the expression of two major proteins involved in the EMT: α-SMA (a mesenchymal marker) and E-cadherin (an epithelial marker). We performed western blot analysis on A549 cells exposed to PMA-Neu or TGF-β, used as a positive control of EMT induction. PMA-Neu induced a significant up-regulation of α-SMA (1.21 ± 0.025) after 24 h compared to control cells (Fig. 2D and F), an effect that was only slightly decreased after 48 h of treatment (1.05 ± 0.102) (Fig. 2E and F). α-SMA upregulation by TGF-β was significant (1.17 ± 0.087) only after 48 h (Fig. 2E and F) compared to control cells. PMA-Neu also downregulated significantly the expression of E-cadherin after 24 h of treatment (0.24 ± 0.101) (Fig. 2D and F), an effect that was amplified at 48 h (0.046 ± 0.032) (Fig. 2E and F). Upon TGF-β administration, E-Cadherin was significant downregulated starting at 24 h (0.253 ± 0.023) (Fig. 2D and F) up to 48 h (0.201 ± 0.131) (Fig. 2E and F).

**Figure 2.**
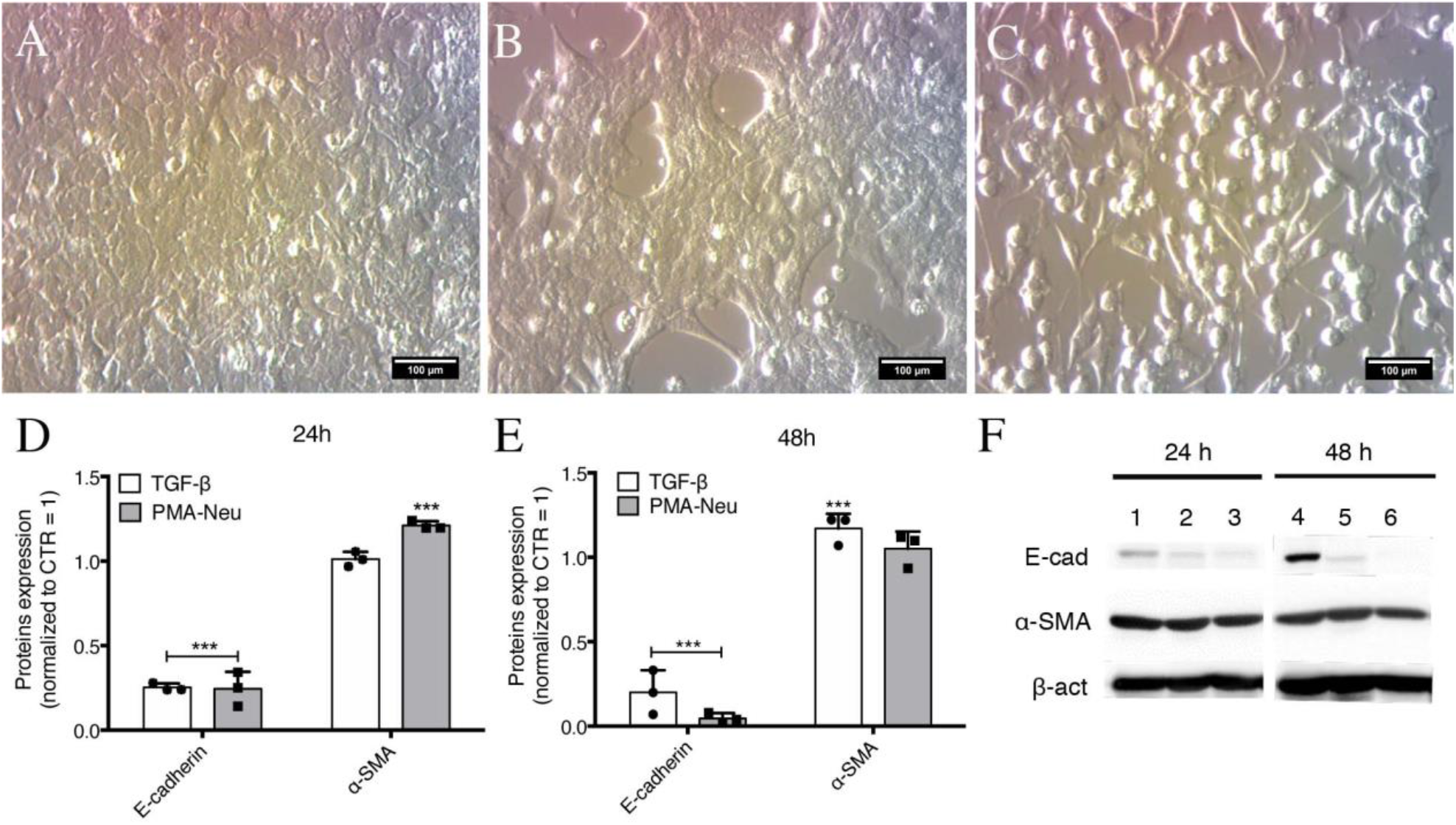
Morphological changes and western blot analysis of A549 treated with PMA-Neu or TGF-β. (A-C) Representative optical images of (A) untreated A549, (B) A549 incubated with TGF-β, or (C) with PMA-Neu after 48 h of treatment. Scale bar = 100 μm. (D,E) Quantification of immunoblots of A549 using anti-E-cadherin or anti-α-SMA incubated with TGF-β or PMA-Neu after (D) 24 h, or (E) 48 h. Data are represented as mean ± SD of three independent replicates. (F) Representative immunoblot of A549 treated with TGF-β or PMA-Neu after 24 and 48 h. Line 1 and 4 = control cells; line 2 and 5 = TGF-β; line 3 and 6 = PMA-Neu. Statistical tests: one-way ANOVA followed by Dunnet test. ***, p<0.001 vs. CTR.

Notably, microscopy of A549 cells treated with PMA-Neu (Fig. 2C) suggests that a portion of cells encounter cell death. The amount of cell death was quantified by propidium iodide (PI) staining 24 h after treatment with PMA-Neu. Flow cytometry analysis confirmed that PMA-Neu induced 33.70 ± 4.37% of cell death compared to control cells (Fig. S1).

Since PMA-Neu were able to induce both the EMT and cell death of A549 cells, we asked if these effects were directly related to NET release. NETs were isolated from either 2.5 × 10^6^ cells (NETs 2.5) or 5 × 10^6^ cells (NETs 5) PMA-Neu. NETs were incubated with A549 cells and EMT induction and cell death were evaluated. Both preparations of NETs induced a significant up-regulation of α-SMA (NETs 2.5 1.35 ± 0.05; NETs 5 2.11 ± 0.12) and decreased E-cadherin expression (NETs 2.5 0.08 ± 0.002; NETs 5 0.35 ± 0.023) after 24 h of treatment (Figure 3A and B), compared to control cells. Interestingly, NETs administration to A549 cells did not induce cell death, in contrast to PMA-Neu (Fig. S1).

**Figure 3.**
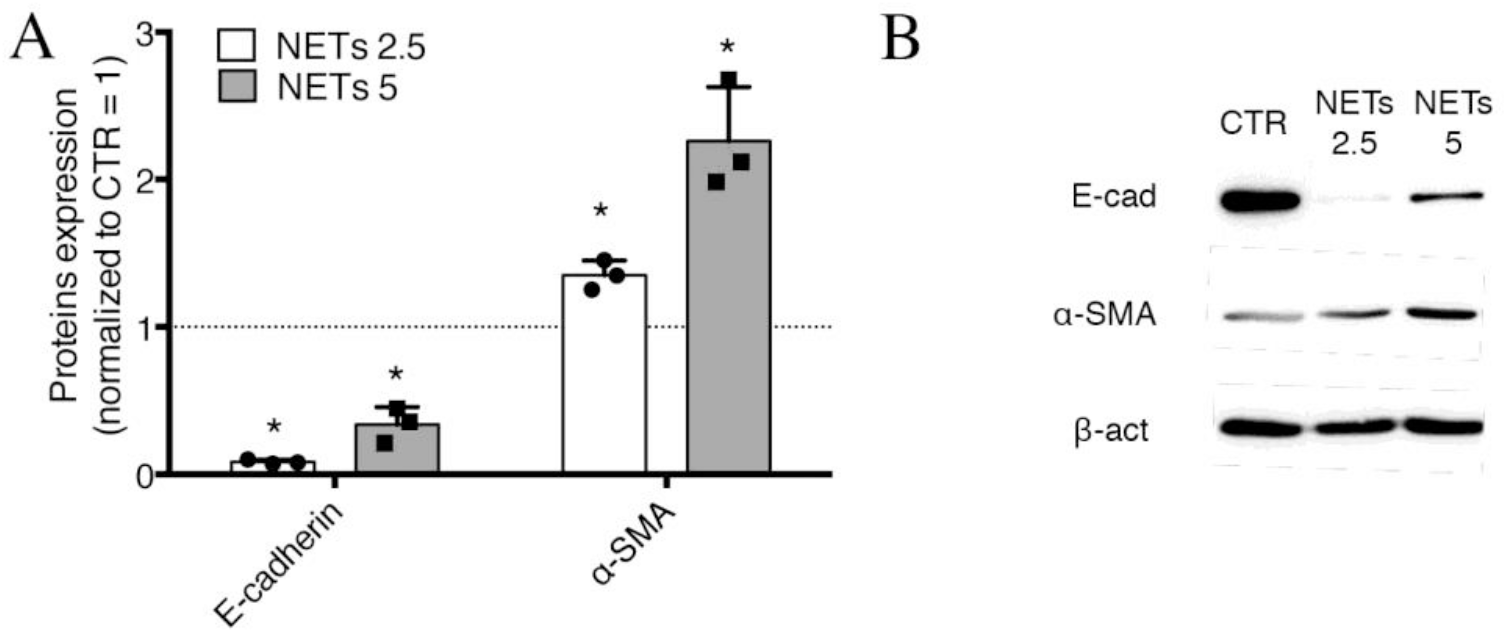
EMT induced by NETs. (A) Quantification of immunoblots using anti-E-cadherin or anti-α-SMA of A549 incubated with NETs 2.5 (derived from 2.5 × 106 PMA-Neu) or NETs 5 (derived from 5 × 10^6^ PMA-Neu) after 24 h of treatment. Data are represented as mean ± SD of three independent replicates. (B) Representative immunoblot of A549 treated with NETs 2.5 or NETs 5 after 24 h. Statistical tests: one-way ANOVA followed by Dunnet test. *, p<0.05 vs. CTR.

### NETs are enriched in the BAL of COVID-19 patients and correlates with neutrophil numbers and IL8 levels

Figure S2 are representative image of immunohistochemical analysis of bronchoalveolar lavage (BAL) of COVID-19 showing that the major cellular components are neutrophils, macrophages, epithelial cells and cellular/nuclear debries. In agreement with recent papers (8, 12, 13), measuring NETs in the BAL of mild (IMW) and severe (ICU) patients, we founded that ICU have a significant higher amount of NETs compared to IMW patients (Fig. 4A). Interestingly, when patients were divided in survivors and non-survivors, we found that the non-survivors group presents more NETs compared to patients who survived (Figure 4B). Finally, we assessed the neutrophil count (Fig. 4C) and the levels of IL8 (Fig. 4D) in the same BAL samples as previously reported (14), and found a significant direct correlation between NETs and neutrophil counts, as well as the levels of IL8.

**Figure 4.**
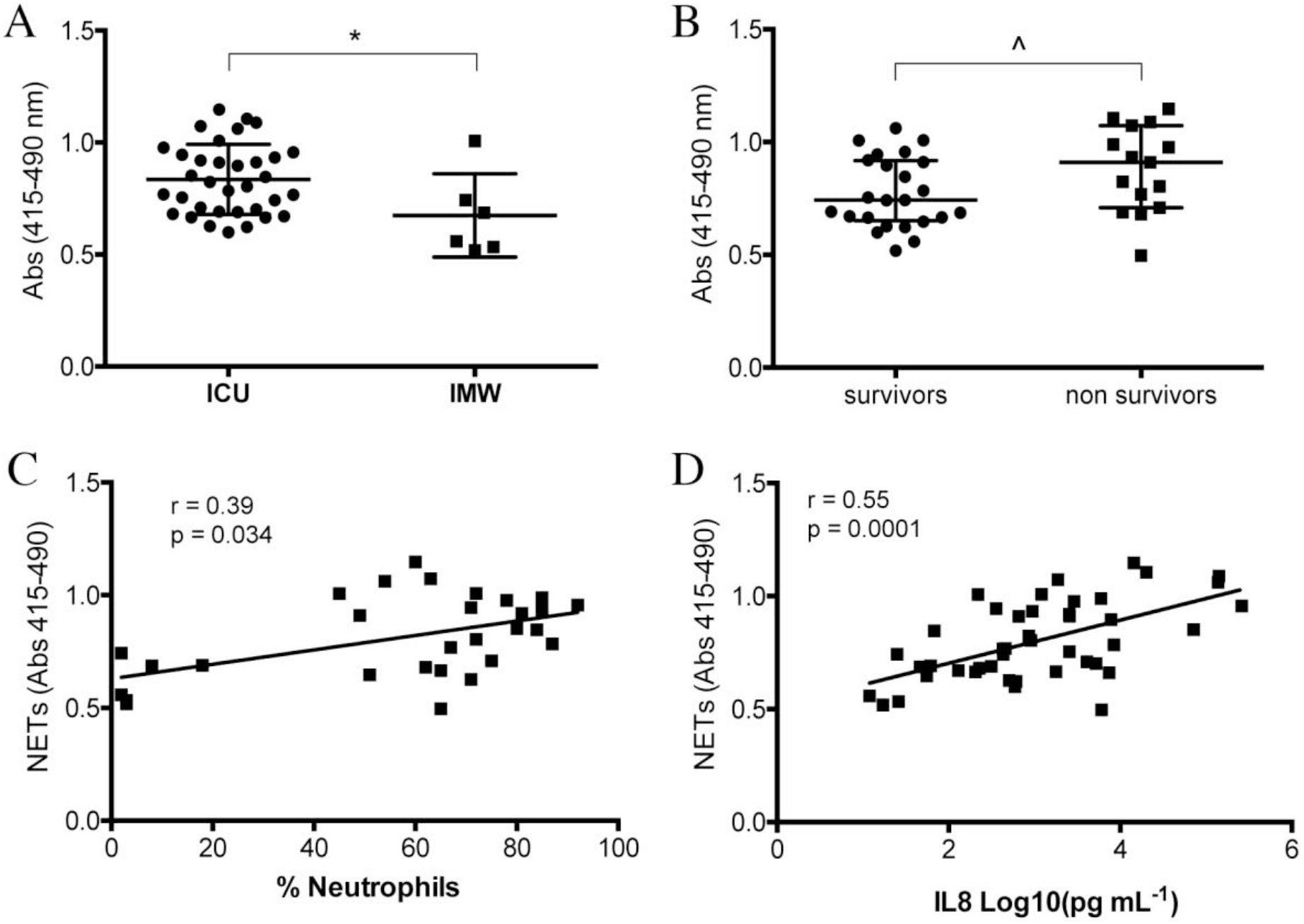
Analysis of BAL COVID-19 patients. (A) Quantification of NETs in BAL of mild (IMW) and severe (ICU) patients. NETs were measured by HNE-DNA complexes quantification subtracting absorbance read at 490 nm to that read at 415 nm. (B) Quantified NETs were compared dividing samples in survivors and non-survivors. Statistical tests: Shapiro-wilk test followed by Mann-Whitney test. (C) Correlation analysis between NETs and percentage of neutrophils counted in the respective BAL sample. (D) Correlation analysis between NETs and IL8 quantified in the respective BAL sample. Correlation analyses were done by Spearman correlation. Data are represented as median (IQR). *, p<0.05; ^, p = 0.05. r = Spearman coefficient.

### The presence of SARS-CoV-2, AMs and neutrophils are necessary for the EMT of lung epithelial cells

We next investigated if in the context of COVID-19, NETs could play a significant role in inducing the EMT. To do so, we set up an alveolar *in vitro* model by culturing A549 cells at the air-liquid interface (ALI). Alveolar macrophages (AM) were seeded at the apical side of A549 cells, while Neu were added to the basolateral chamber of the transwell (15, 16). Our *in vitro* alveolar model was, then, inoculated with SARS-CoV-2 (Neu+AM+SARS-CoV2). As a control, A549 cells were cultured alone, or with SARS-CoV2, or with Neu+SARS-CoV2. After 48 h of incubation, we observed that only in the wells containing Neu+AM+SARS-CoV2 epithelial cells encountered the EMT, as demonstrated by a significant up-regulation of α-SMA (1.6 ± 0.321), together with a down-regulation of E-cadherin (0.723 ± 0.059), compared to control cells (Fig. 5A and B). When epithelial cells were treated with SARS-CoV2 only, the expression of E-cadherin was significantly reduced (0.667 ± 0.249) but α-SMA levels were not significantly increased, while Neu+SARS-CoV2 cultures induced the up-regulation of α-SMA (1.339 ± 0.216), but not the downregulation of the E-cadherin (Fig. 5A and B). Altogether these data suggest that the presence of AM, Neu and the virus are necessary for an efficient EMT of lung epithelial cells.

**Figure 5.**
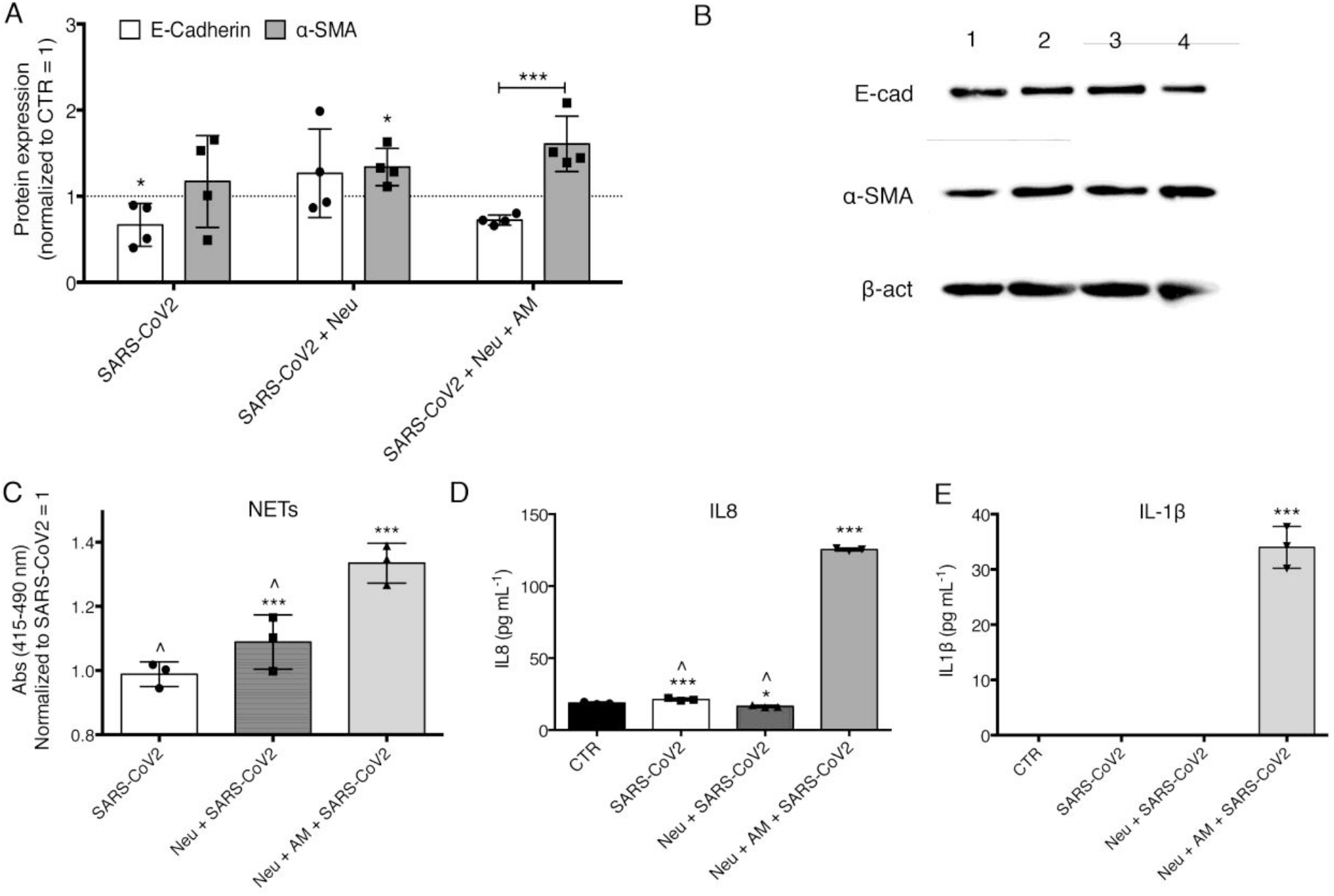
Western blot of EMT NETs and NETs/cytokines analysis in alveolar *in vitro* model. (A) Semiquantitative analysis of four independent immunoblots and (B) representative immunoblot using anti-E-cadherin or anti-α-SMA or β-actin of A549 co-cultured with SARS-CoV2, Neu+SARS-CoV2, or Neu+AM+SARS-CoV2 after 48 h of treatment. Line 1 = control cells; line 2 = SARS-CoV2; line 3 = Neu+SARS-CoV2; line 4 = Neu+AM+SARS-CoV2. Statistical tests: one-way ANOVA followed by Dunnet test. (C) NETs quantification by DNA-HNE complexes measurement. (D) IL8 and (E) IL1β quantification by ELISA assay. Statistical tests: Shapiro-wilk test followed by one-way ANOVA and Dunnet test. Data are represented as mean ± SD. ***, p<0.001 vs. CTR; *, p<0.05 vs. CTR; ^, p<0.001 vs. Neu+AM+SARS-CoV2.

To clarify if NETs are produced by Neu and involved in this *in vitro* experimental setting, we measured HNE-DNA complexes in culture media after 48 h of incubation, but also IL8 and IL1β, two most known effectors of NETosis. In agreement with the role played by NETs in the induction of the EMT, we found a significant higher induction of NETs in Neu+AM+SARS-CoV2 compared to Neu+SARS-CoV2 and, as expected, SARS-CoV2 alone (Fig. 5C). Similarly, the quantification of IL8 (Fig. 5D) and IL1β (Fig. 5E) showed that these two cytokines are only produced in Neu+AM+SARS-CoV2 cultures, in agreement with the well-known capacity of AM to secrete these cytokines (17).

### EMT marker in lung biopsies in COVID-19 dead patients

Histopathological examination of lung tissues obtained from two patients who died of COVID-19 pneumonia demonstrated exudative and proliferative features of diffuse alveolar damage, corresponding to an early/intermediate phase of the disease (18). Immunohistochemical analyses showed a subset of pneumocytes expressing both epithelial and mesenchymal markers (cytokeratin 7 and α-SMA) (Fig. 6), thus confirming that these epithelial cells acquired a mesenchymal phenotype via an EMT process.

**Figure 6.**
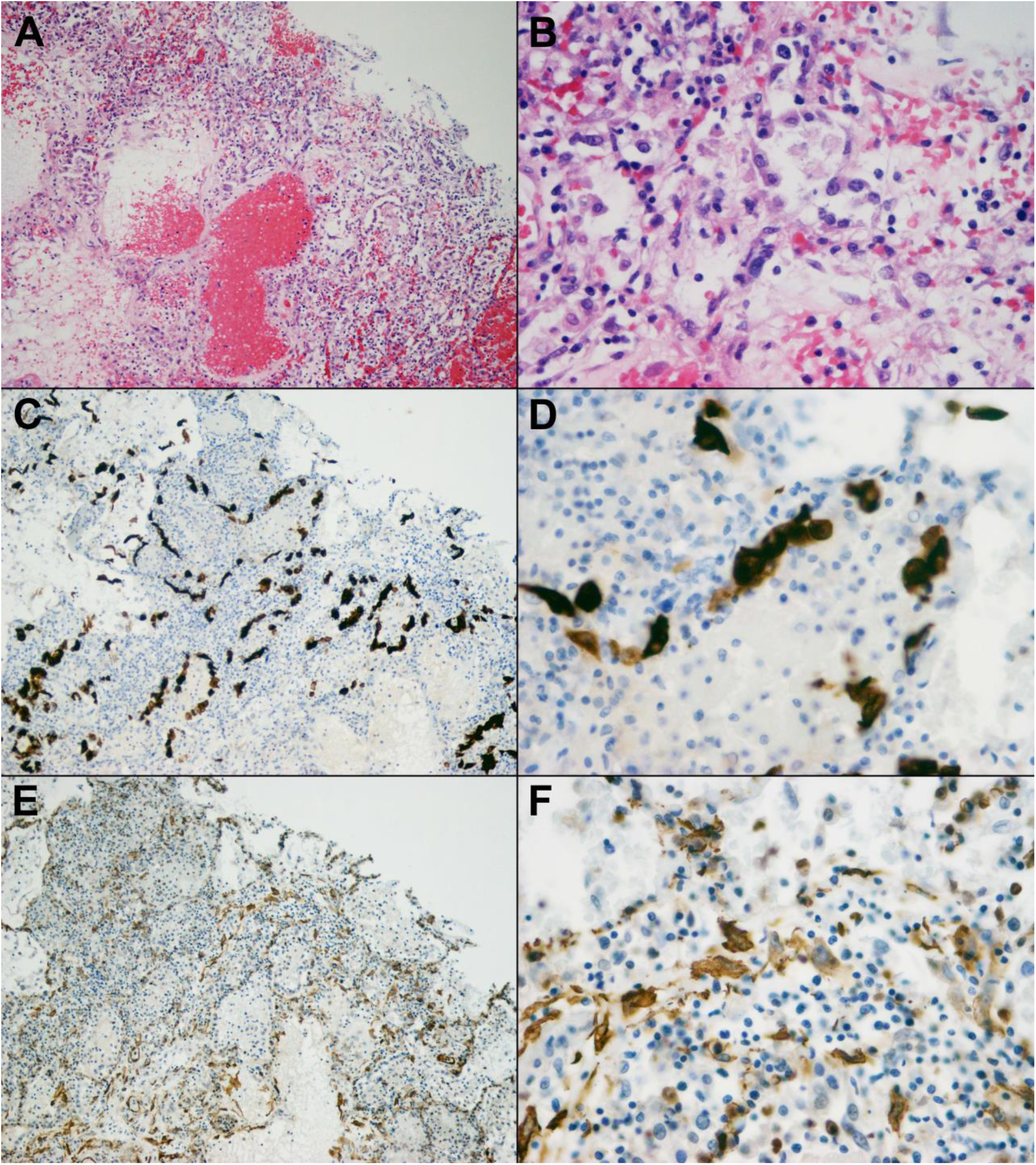
Lung tissue of COVID-19 deceased patient with DAD in proliferative phase. (A-B) Vascular congestion, oedema, hyperplastic pneumocytes, a few granulocytes, fibrin deposition and focal hyaline membranes in alveolar spaces; H&E OM (A) 10x and (B) 40x. (C-D) anti-Cytokeratin 7 immunohistochemistry staining residual epithelial cells; immunohistiochemistry, OM (C) 10x and (D) 40x. (E-F) anti-α-SMA immunohistochemistry staining of mesenchymal stromal and vascular cells; OM (E) 10x and (F) 40x. (F) The same epithelial cells of (D) express α-SMA indicating the gain of a mesenchymal phenotype.

## Discussion

Our study demonstrated for the first time that NETs are sufficient to induce EMT in the A549 alveolar epithelial cell line. Moreover, we confirmed a significant correlation between NET induction in the lung and the severity of COVID-19. We also demonstrated in an *in vitro* model that NETs, possibly sustained by the secretion of pro-inflammatory cytokines from AMs, are involved in the inducement of EMT. Notably, lung tissues of COVID-19 deceased patients showed that epithelial cells are characterized by increased mesenchymal markers.

We found that neutrophils efficiently induce the EMT via NETosis, an observation that, at the best of our knowledge, has never been demonstrated before in the context of lung fibrosis and, more importantly, in severe COVID-19. NET-related injuries are usually studied in the context of the endothelium (19, 20). In particular, the prothrombotic role (21) of NETs and their ability to drive endothelial-mesenchymal transition (22) has been thoroughly analyzed. However, recent reports suggested that the infiltration of neutrophils in the alveolar space interferes with cell-cell adhesion of lung epithelial cells through HNE (23), also, that neutrophils induce the EMT by releasing TGF-β, neutrophil gelatinase-associated lipocalin (NGAL) (24) or Proteinase-activated receptor 4 (PAR4) (25). Herein, we demonstrate that NETs can also directly activate the EMT program in A549 cells, a commonly used model of type II pneumocytes. We showed that the addition of purified NETs to A549 cells induced a significant overexpression of the mesenchymal marker α-SMA after 24 h of treatment, together with a decrease of E-cadherin expression (Fig. 3). Comparing PMA-Neu and NETs effect, we found that although both treatments are able to trigger the EMT (Fig. 2 and 3), the presence of neutrophils is uniquely able induce the death of A549 cells (Fig. S1). These results not only confirm data already present in the literature, but also shed new light on the mechanism that leads to the EMT in the lung: exaggerate NETosis not only induces tissue damage (1, 6) but it also triggers the EMT in lung epithelial cells.

Another important finding of our study is the presence of NETs in the BAL of COVID-19 patients. Given the high percentage of NETs in the peripheral blood and the exaggerate infiltration of neutrophils in alveolar spaces of SARS-CoV-2-infected patients (26, 27), the role of these phagocytes in the pathogenesis of COVID-19 was clear since the very beginning. NET levels in different samples of SARS-CoV2-infected patients significantly correlated with the disease severity, suggesting that NETosis may play a relevant role in COVID-19 (7, 8, 12). Here, we confirmed and expanded previous observations by quantifying NETs in the alveolar micro-environment. We took into consideration two groups of patients, those with mild disease (patients that were admitted to the hospital with signs and symptoms of bilateral interstitial pneumonia but did not required intubation) and those with severe disease (who needed ICU support). Our data demonstrated higher levels of NETs in severe patients compared to IMW (Fig. 4A). In addition, NET levels correlated to the percentage of neutrophils in the BAL (Fig. 4C), as well as with the levels of IL8 (Fig. 4D) and the disease severity (Fig. 4B).

Because of the high levels of NETs in the BAL of COVID-19 patients and their potential role in inducing the EMT, we next focused on the correlation between NETs and the EMT by setting up an *in vitro* alveolar model that we infected with SARS-CoV-2. To mimic the alveolar microenvironment A549 were cultivated at the ALI to have the contact with air, such as under physiologic conditions. Next, we added AM onto the apical side of A549 and Neu in the basolateral chamber of the transwell. SARS-CoV2 was inoculated on the apical side and after 48 h only the Neu+AM+SARS-CoV2 culture significantly induced the EMT in A549, in contrast to Neu+SARS-CoV2 and SARS-CoV2 alone conditions (Fig. 5A and B). Another recent study suggested that SARS-CoV2 infection may trigger EMT-like molecular changes in A549, as determined by the upregulation of the *ZEB1* gene together with the downregulation of *EPCAM* (28). However, in our experimental model, we demonstrated that the sole presence of SARS-CoV2 is not sufficient to induce an efficient EMT, as measured by α-SMA/E-cadherin regulation (Fig. 5A). Notably, also in the case of Neu+SARS-CoV2 we did not see a strong EMT, as measured by α-SMA/E-cadherin protein modulation in A549 (Fig. 5A), although Neu were stimulated to secrete NETs by SARS-CoV2 (Fig. 5C), as previously described by Veras et al (13). Thus, we hypothesize that the amount of NETs released by Neu in Neu+SARS-CoV2 is not sufficient to elicit the EMT in A549 cells. In fact, when we added AM, we observed an enhancement of NETs release probably due to the capacity of these cells to secrete IL8 and IL1β (Fig. 5D and E), two proteins known to efficiently drive NETosis (29). We are aware about the low endogenous expression by A549 of ACE2, the specific receptor for SARS-CoV2 cell entry (30, 31). However, our results are even more interesting because emphasizes the critical role of innate immunity in SARS-CoV2 infection despite the internalization of virus by cells. In particular, we would like to propose a model in which macrophages, by releasing IL8 and IL1β, two cytokines significantly increased in the plasma (32, 33) and BAL-fluid of severe COVID-19 patients (14), potentiate the capacity of NETs released by Neu to amplify the noxious activity on lung cells, favoring the EMT. Our data are in agreement with previous studies that showed the existence of a feedback loop between neutrophils/NETs and release of IL8 (29) and IL1β (5).

The immunohistochemistry analysis of lung tissue obtained *post-mortem* from deceased COVID-19 patients supported our *in vitro* findings, showing a subset of pneumocytes expressing mesenchymal markers (Fig. 6), thus confirming that epithelial cells acquired a mesenchymal phenotype.

Futures studies will be focused on

In conclusion, this study highlights the contribution of neutrophil activation and NET generation in the induction of the EMT. Several inflammatory disorders besides ARDS, such as autoimmune diseases and chronic lung graft rejection have been associated at various degrees with alveolar and small airway influx of Neu and with their activation (34, 35). We, thus, hypothesize that NETosis-induced EMT should be considered as an important pathogenic step of lung fibrosis consequent to neutrophilic inflammation. Our findings also suggest that future therapeutic interventions could target this mechanism.

## Methods

### COVID-19 patients

BAL of 28 adults positive for SARS-CoV-2 infection, diagnosed by real-time PCR on nasopharyngeal swab, were collected. N = 25 were admitted to the Intensive Care Unit (ICU) at the Luigi Sacco Hospital (Milan, Italy); N = 6 were admitted to the Intermediate Medicine ward (IMW) of the IRCCS Policlinico San Matteo Foundation. BALs were collected as previously described (14). After centrifugation, supernatants were analyzed to quantify IL8 release; cell pellets were fixed, and neutrophils were counted after staining with Papanicolaou.

Additional BAL samples were fixed in 10% buffered formalin and cell-blocks were done; three-μm paraffin sections were stained by H&E for cytological examination.

Lung autopsy tissues from 2 patients died of SARS-CoV-2 pneumonia were fixed in 10% buffered formalin for 48 h. Three-μm paraffin sections were stained by H&E. Immunohistochemistry reactions were performed on the most representative area by using anti-Cytokeratin 7 (clone SP52, Ventana) and anti-α-SMA (clone 1A4, Ventana).

### Immune cells isolation

Neutrophils were isolated from peripheral blood of healthy donors. Blood was stratified by Lympholyte® Cell Separation Media (EuroClone, Milan, Italy) and after centrifugation at 450 g for 30 min mononuclear cell phase was eliminated to allow neutrophils isolation. A solution of 3% Dextran (200 kDa) in 0.85% NaCl was added to the remaining phases of neutrophils and erythrocytes followed by a 30 min incubation at room temperature, allowing the precipitation of erythrocytes. Supernatant was collected and washed with PBS (EuroClone). After centrifugation at 450 g for 10 min, pellet was resuspended with VersaLyse Lysing Solution (Beckman Coulter S.r.l., Milano) for 20 min at room temperature in the dark. After the addition of PBS, sample was centrifuged again at 450 g for 10 min. This passage was repeated until the pellet was found to be free of erythrocytes.

Alveolar macrophages (AM) were isolated from BAL of four patients without lung infection and with a cytologic count showing AM > 76% and lymphocytes < 14%. BAL were filtered with a sterile gauze, centrifuged at 450 g for 10 min and then pellet of cells were cultured in suspension for 24 h in DMEM containing 10% FCS, P/S and L-glutamine.

### NETs isolation

To induce the release of NETs and to isolate them we followed a published protocol with some modifications (36). 5 × 10^6^ isolated neutrophils were cultivated in 60 cm^2^ plate with the addition of 100 nM PMA (Sigma-Aldrich S.r.l.,Milan, Italy) for 4 h at 37°C. After that, supernatant was discarded, and the layer of NETs present onto the bottom of the plate was harvested using PBS and collected into falcon. After centrifugation at 450 g for 10 min, cell-free NETs-rich supernatant was divided in 1.5 mL Eppendorf and centrifuged at 18,000 g for 10 min at 4°C.

### Confocal microscopy of neutrophils and NETosis

2 × 10^6^ of isolated neutrophils were cultivated onto coverslips and incubated with 100 nM PMA (PMA-Neu). After incubation, cells were washed with PBS and fixed with 4% of paraformaldehyde for 10 min. After three washes with PBS, cells were treated with blocking solution (1% BSA in PBS) for 1 h at room temperature. Cell were than stained with anti-H3 polyAb (dilution 1:100 – PA5-31954 - Invitrogen - Life Technologies) or anti-MPO polyAb (diluition 1:50 – DOM0001G - Invitrogen) or anti-HNE mAb (diluition 1:50 – MA1-40220 - Invitrogen) in 0.5% BSA in PBS for 1 h at room temperature. After three washes in PBS, secondary mAbs (anti-mouse and anti-rabbit IgG H&L, Alexa Fluor® 488 - Abcam; anti-rabbit IgG H&L, Alexa Fluor® 647 - Invitrogen) was added in 0.5% BSA in PBS for 1 h at room temperature. Neutrophils without treatment with PMA were used as control. Coverslips were than mounted using ProLong™ Gold Antifade Mountant with DAPI (Invitrogen) and analyzed with confocal microscopy (Fluoview FV10i, Olympus).

### SARS-CoV2

A wild strain of SARS-CoV-2 was isolated in VERO E6 cell line from a nasal swab of a COVID-19 infected patient. Virus was propagated and titrated to prepare a stock to be stored at −80°C to be used for all the experiments at a concentration of 100 TCID50.

### Cell culture

A549 cell line (purchased from ATCC^®^) was cultivated with DMEM high glucose supplemented with 10% of FBS, 1% of penicillin-streptomycin solution and 1% of L-glutamine (all purchased by EuroClone). Cell were cultivated at 37°C with 5% of CO_2_ and harvested when they reached the 80% of confluence.

### EMT induction (A549 + neutrophils/NETs)

To study the induction of EMT on A549 by PMA-Neu/NETs, 0.2 × 10^6^ A549 were cultured on 12-well plate for 24 h. 2.5 × 10^6^ and 5 × 10^6^ PMA-Neu /NETs were added in each well and after 24 and 48 h supernatants were discarded and A549 were lysed to extract all proteins to perform western blot analysis. As positive control we treated A549 with 10 μg mL^−1^ of TGF-β.

Cell death was evaluated labeling A549 with PI after 24 h of incubation with 5 × 10^6^ PMA-Neu /NETs and analyzed with flow cytometry.

### Alveolar *in vitro* model

To mimic the alveolar space *in vitro*, 0.5 × 10^6^ A549 cells were cultured onto transwell inserts (0.4 μm-pore size - Corning Costar) in ALI to allow a good polarization of cells. After 14 days, different conditions were prepared:

Only A549 as control
A549 + SARS-CoV2 (added onto A549 in 20 μL to not modify the polarization of cells)
A549 + 2.5 × 10^6^ neutrophils + SARS-CoV2
A549 + 0.5 × 10^6^ AM + 2.5 × 10^6^ neutrophils + SARS-CoV2

After 24 and 48 h, medium in the lower compartment was collected to perform cytokines quantifications, transwell inserts were cut around the membrane edges to lyse cells to extract all proteins and to perform western blot analysis.

### Western blot

Cells were washed with PBS, lysed with lysis buffer (50 mM Tris-HCl [pH 7.4], 150 mM NaCl, 10% glycerol, 1% NP-40, protease inhibitor cocktail (Sigma Aldrich) and phosphatase inhibitor (Roche), gently vortexed for 20 min at 4°C and centrifuged for 15 min at 13,200 rpm at 4 °C. Supernatants were quantified by Pierce™ BCA Protein Assay Kit (Thermo Fisher Scientific).

Twenty micrograms of proteins from A549 cell extracts were loaded and separated in 8% SDS-PAGE. After electrophoresis, the gels were transferred to polyvinylidene difluoride membranes (Millipore), therefore blocked (5% no fat milk in 0.1% Tween 20 TBS) and incubated with the primary Ab (1:1000 in TBST + 2% BSA; overnight at 4 °C or 2 h at room temperature): anti-E-Cadherin [M168] (ab76055, Abcam), anti-α-SMA [E184] (ab32575, Abcam), and anti-β-Actin (MAB1501R, Chemicon). After wash, the membranes were incubated with the appropriate horseradish-peroxidase conjugated secondary Ab (1:5000 in TBST + 2% BSA; 2 h at room temperature; anti-mouse A4416 and anti-rabbit A0545, Sigma). The immunoreactivity was detected by ECL reagents (Amersham), acquired with the ChemiDoc imaging system (Image Lab, Bio-Rad).

### Cytokines and NETs quantification

To quantify cytokines released in the alveolar *in vitro* model, ELISA assays were performed. We quantified IL8 with SimpleStep ELISA® Kit (Abcam) and IL1β with human IL-1β/IL-1F2 Immunoassay (R&D Systems) was titered using a commercial enzyme-linked immunosorbent assay kit (Human IL-1β/IL-1F2 Immunoassay, R&D Systems) following the manufacturer’s instructions and the results were expressed as pg ml^−1^. All determinations were measure in same session. For IL8 quantification in BAL of COVID-19 patients we referred to already published quantifications (14). To quantify NETs we measured HNE-DNA complexes using ELISA specific for HNE and anti-DNA Ab of Cell Death Detection ELISA^PLUS^ (Roche). Briefly, 100 μL of sample (diluted 1:100) was added to anti-HNE ELISA kit (Thermo Fisher Scientific) for 1 h at room temperature. After three washes, anti-DNA Ab was added on each well followed by an incubation of 2 h at room temperature. After three washes, substrate was added for 1 min. After adding stop solution, absorbance was read at 415 nm and 490 nm. The results are represented as the result of this equation:

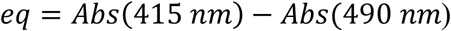

### Statistical analysis

*In vitro* results presented in this study were analyzed by one-way ANOVA followed by Dunnet test. Results were obtained from three/four independent experimental replicates represented as mean ± SD. BAL analyses were done by Mann-Whitney test, while correlation analyses were done calculating Spearman coefficient. Data are represented as median (interquartile range – IQR). All analyses were carried out with a GraphPad Prism 6.0 statistical program (GraphPad software, San Diego, CA, USA). A value p < 0.05 was considered statistically significant.

### Study approval

Research and data collection protocols were approved by the Institutional Review Boards (Comitato Etico di Area 1) (prot. 20100005334) and by IRCCS Policlinico San Matteo Foundation Hospital (prot.20200046007). Written informed consent was obtained by all conscious patients. For unconscious patients the inform consent was obtained after recovery and for non-survivors was waived in accordance to the Italian law.

## Author’s contribution

PL, MF, LD designed the research; PL, FV, BS, PE, GE performed the *in vitro* experiments; D’AM, MM, BS, FV, PL handled BAL samples; FT, CR collected BAL from COVID-19 patients; VBM, DeLA performed western blot analysis; DeAM quantified IL1beta; LG, NM, CL performed immunohistochemistry analyses on COVID-19 biopsies; PL, FV, analyzed data; PL, BS, FV, MF wrote the main text; ALL AUTHORS revised the manuscript.

## Acknowledgments

Prof. Zanoni Ivan (Division of Immunology, Division of Gastroenterology, Boston Children's Hospital, Harvard Medical School, Boston, MA, United States) to revise the manuscript.

Monti Manuela PhD (Laboratory of Biotechnology, Center of Regenerative Medicine Research, IRCCS San Matteo Foundation, Pavia, Italy) for the use of confocal microscopy.

Testa Giorgia (Pediatrics Clinic, IRCCS Policlinico S. Matteo Foundation, Pavia, University of Pavia Italy) for the analysis of IL1β.

Fondazione Cariplo (COVIM project); Ministry of Health funds to IRCCS Foundation Policlinico San Matteo Grant (Ricerca Corrente) and Ministry of Health funds COVID-2020-12371760. The funders had no role in study design, data collection and analysis, or preparation of the manuscript.

