## supplementary materials for "Neutrophil extracellular traps induce the epithelial-mesenchymal transition: implications in post-COVID-19 fibrosis"

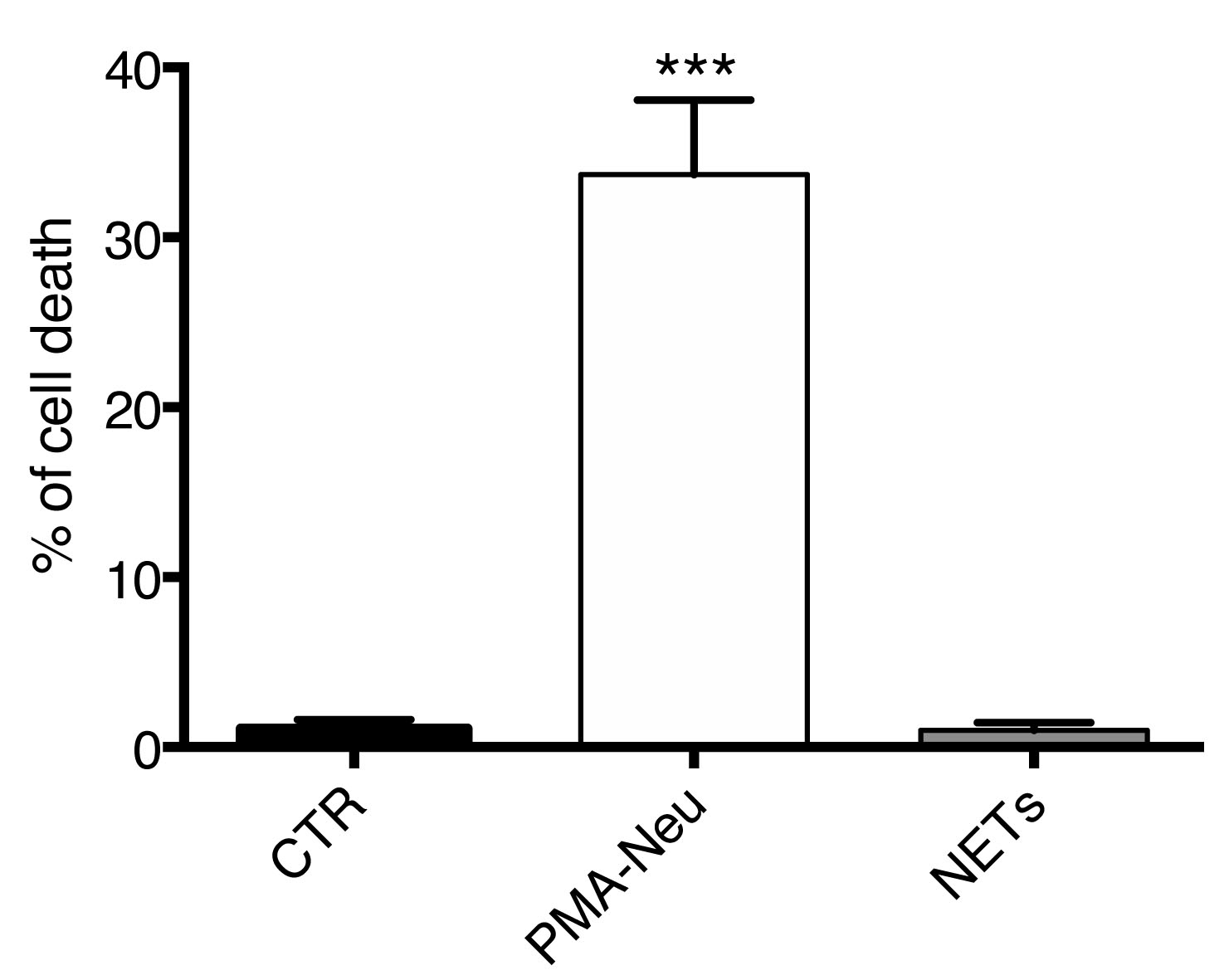

Figure S1. Cell death analysis. Cell death of A549 incubated with 5 x 10^6^ PMA-Neu or NETs (24 h) was evaluated by flow cytometry labeling A549 with PI before the acquisition. Data are represented as mean ± SD of three independent replicates. ***, p<0.001 vs. CTR.

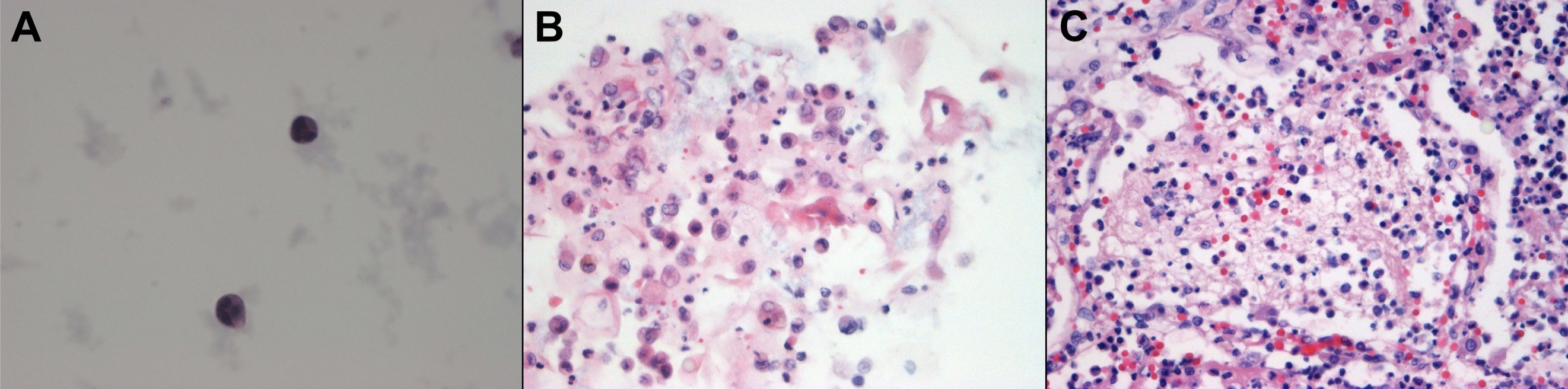

Figure S2. Morphological evaluation of BAL samples. (A) Two neutrophils in in the cytological smear; OM 100x. (B) Cell block characterized by macrophages, epithelial cells, neutrophils and cellular/nuclear debries; H&E OM 40x. (C) histological evaluation of lung tissue with macrophages, epithelial cells, neutrophils and cellular/nuclear debries in alveolar spaces; H&E OM 40x.
